# Single-nucleus transcriptomics reveal disrupted pathways in the prefrontal cortex of *Scn2a*-deficient mice

**DOI:** 10.1101/2025.09.16.676420

**Authors:** Ye-Eun Yoo, Purba Mandal, Zhengkuan Tang, Zaiyang Zhang, Jingliang Zhang, Xiaoling Chen, Morgan Robinson, Muriel Eaton, Brody Deming, Manasi Halurkar, Harish Kothandaraman, Luke C. Dabin, Boyu Jiang, Hongyu Gao, Chongli Yuan, Nadia Lanman, Yunlong Liu, Jungsu Kim, Priyanka Baloni, Yang Yang

## Abstract

Truncating variants in *SCN2A*, which encodes the NaV1.2 sodium channel critical for action potential initiation and propagation, are associated with autism spectrum disorder (ASD) and epilepsy. To investigate *SCN2A* deficiency–related phenotypes, we developed a preclinical mouse model with under 50% NaV1.2 expression, exhibiting neuronal hyperexcitability and social deficits. However, the neuronal populations and molecular alterations underlying these phenotypes at single-cell resolution have not been investigated. In this study, we conducted single-nucleus RNA sequencing (snRNA-seq) of wild-type (WT), homozygous *Scn2a*-deficient (HOM) mice, and HOM mice with *Scn2a* restoration (HOM-FlpO) to examine the effects of *Scn2a* level on gene expression in the medial prefrontal cortex (mPFC), a critical brain region related to ASD development. Differential expression analysis in GABAergic and glutamatergic neurons between genotypes revealed gene expression enriched in neurotransmitter activity regulation and synapse organization. Lastly, snRNA-seq results in HOM-FlpO identified genes that were rescued after *Scn2a* restoration. These results reveal that reduced *Scn2a* expression disrupts RNA transcriptomes in multiple neuronal subtypes, providing insight into cell type–specific mechanisms underlying *SCN2A*-related disorders.

## Introduction

The *SCN2A* gene, encoding voltage-gated sodium channels NaV1.2, has a critical role in regulating neuronal excitability (1–3). It is mainly expressed in the axon initial segment (AIS) and dendrites of glutamatergic and GABAergic neurons in the central nervous system, where it contributes to the generation and propagation of actional potential (4–6). Mutations in sodium channels can disrupt neuronal excitability, leading to atypical neurodevelopment, impaired cognitive function, and epileptic seizures (7–9). Several genetic studies have identified *SCN2A* as one of the most prominent risk genes for autism spectrum disorders (ASDs) (10–13). *SCN2A* genetic variants have been repeatedly identified in patients with autism, epilepsy, and developmental delays (2, 14, 15).

In order to study *Scn2a* deficiency in a preclinical mouse model, we previously developed and characterized *Scn2a* gene trap mice (16). To address gene expression changes caused by *SCN2A* deficiency, we conducted bulk RNA sequencing (RNA-seq) in the medial prefrontal cortex (mPFC) (17). The mPFC is an important brain region involved in higher cognitive functions, such as social interaction, and has been implicated in autism and developmental delays (18, 19). A myriad of neuronal cell types in the mPFC integrate inputs from multiple brain regions (20–23). Comprehensive spatial transcriptomic analyses further reported a diversity of cellular populations, including immature GABAergic neurons in the adult mouse mPFC, characterized by markers associated with developmental plasticity (24, 25). These immature neurons are thought to contribute to cortical circuit remodeling and functional adaptability (26). Proper cell-type composition in the mPFC is essential for normal cognitive function, and its disruption is implicated in multiple neurodevelopmental and neuropsychiatric disorders (27–29).

In prior work, we and others have demonstrated that pyramidal neurons in the mPFC of adult *Scn2a*-deficient mice exhibit hyperexcitability, which was normalized by restoring *Scn2a* expression (17). Interestingly, we also found that social deficits in *Scn2a*-deficient mice were rescued by adult *Scn2a* restoration (30). However, bulk RNA sequencing only captures average gene expression across heterogeneous cell populations, limiting the ability to detect changes in specific cell types. It remains unclear which neural cell types are most affected by *Scn2a* deficiency and how changes in gene expression contribute to cellular dysfunction and behavioral phenotypes. Characterizing cell–type–specific molecular effects of Scn2a deficiency and their modulation by genetic restoration is critical for elucidating the mechanisms of *SCN2A*-related disorders and identifying therapeutic targets.

To investigate cell-type-specific molecular and cellular changes induced by *Scn2a* deficiency, we performed single-nucleus RNA sequencing (snRNA-seq) in the mPFC of the WT mice, homozygous *Scn2a*-deficient mice, and Hom mice with restored *Scn2a* (WT, HOM, HOM-FlpO, hereafter). We observed an unexpected increase in immature GABAergic neurons in HOM. Differentially expressed genes (DEGs) between HOM and WT were significantly enriched in synaptic transmission and cell signaling-related pathways. Additionally, cell-cell communication network analysis revealed that increased intercellular communication in HOM mPFC was ameliorated by FlpO-induced *Scn2a* restoration. This study provides the framework for identifying the molecular basis for potential treatments by analyzing transcriptomic differences between WT and *Scn2a*- deficient mice at the single-cell level.

## Methods

### Animal model

C57BL/6N-*Scn2a1^tm1aNarl^*/Narl (*Scn2a^+/gt^*) mice were generated and characterized as previously described (16). Briefly, heterozygous (HET, *Scn2a^+/gt^*) mice were bred to obtain homozygous (HOM, *Scn2a^gt/gt^*) and wild-type (WT, *Scn2a^+/+^*) littermates for experiments. Mice were group housed (3-5 mice per cage) on vented cage racks with a 12-hour light/dark cycle. Food and water were given *ad libitum*. Animals were tested under guidelines provided by Purdue Institutional Animal Care and Use Committee and the National Institutes of Health (NIH).

### Adeno-associated virus (AAV) transduction

For *Scn2a* HOM-FlpO group, AAV-PHP.eB vectors were prepared and systemically delivered to mice as previously described (17). For *Scn2a* HOM-FlpO group, AAV-PHP.eB vectors were prepared and systemically delivered to mice as previously described (17). 3-5-month-old mice received 2X10^11^ viral particles of AAV-FlpO (PHP.eB-EF1a-mCherry-IRES-Flpo-WPRE-hGH) or control AAV-mCherry virus (PHP.eB-Ef1a-DO-mCherry-WPRE-pA) via tail vein injection.

### Three-chamber test

Three months after AAV-FlpO injection, a three-chamber sociability test was performed to test social behavior. Sociability tests involved placing mice to habituate in a three-chamber apparatus. A novel WT mouse was placed in the holding cylinder of one chamber as a social target, and the other cylinder of the chamber was left empty. The subject mice were allowed to explore the chambers for 10 minutes. Interaction time was defined as the time spent sniffing an empty cylinder or a cylinder with a social target.

### Library preparation and single-nucleus RNA sequencing (snRNA-seq)

The frozen prefrontal cortex (PFC) brain tissues of four replicates, including two females and two males per group (WT, HOM, HOM-FlpO), were used. Mice were deeply anesthetized with ketamine/xylazine (100/10 mg/kg, intraperitoneal injection), and brains were rapidly extracted and flash-frozen in liquid nitrogen. Coronal brain sections containing the medial prefrontal cortex (mPFC) were prepared, and the mPFC regions were dissected using a cold biopsy punch as previously described (31). The tissue samples were homogenized in the chilled lysis buffer (10 mM Tris-HCl, pH 7.5, 10 mM NaCl, 3 mM MgCl2, 0.1% Nonidet P40 Substitute, 1 mM DTT, 0.2 U/µl RNase inhibitor) using a Dounce homogenizer. Lysates were centrifuged and filtered with a 40 µm cell strainer. Dissociated nuclei were resuspended in PBS containing BSA and RNase inhibitor, and then purified by cell sorter for downstream library preparation. snRNA-seq libraries were generated using Chromium Next GEM Single Cell 3ʹ Reagent Kits v3.1 (10X Genomics), following the manufacturer’s instructions. Approximately 6,000 nuclei per sample were targeted for capture on the Chromium platform. Final libraries were sequenced on an Illumina NovaSeq 6000, with a targeted sequencing depth of ∼60,000 reads per nucleus.

### Processing of single-nucleus RNA sequencing data

The sequencing data were processed via the standard pipeline of the Cell Ranger single-cell software suite 6.1.1 (10x Genomics). The FASTQ files generated from the pipeline were aligned to the mouse reference genome (mm10) and used to create the gene expression matrix based on the count of unique molecular identifiers (UMIs) assigned to each transcript by Cell Ranger counts. The filtered gene expression matrices were used for subsequent downstream analysis.

### Single-nucleus RNA sequencing data analysis

The gene expression matrices were further analyzed using the Scanpy package (32) (v1.10.2). To remove low-quality nuclei, cells that have fewer than 500 unique genes and more than 6000 genes were removed, and to exclude cytoplasmic transcripts and avoid biases introduced during nuclei isolation and ultracentrifugation, cells with greater than 10% of reads mapping to mitochondrial genes were removed. The dataset was further filtered for doublets using Scrublet package (33) (v0.2.3) using the default parameters. A total of 47,291 cells were retained for downstream analysis, including 14,204 cells from WT, 15,097 cells from HOM, and 17,990 cells from HOM-FlpO.

The filtered data were normalized by total counts per cell using the ‘scanpy.pp.normalize_per_cell’ function, followed by log-transformation with the ‘scanpy.pp.log1p’ function (natural log, log(x+1) transformation applied to the data matrix). To address potential batch effects across samples, we selected the top 3,000 highly variable genes using ‘scanpy.pp.highly_variable_genes’ and used them for principal component analysis (PCA, 50 PCs) and batch-balanced k-nearest neighbors (BBKNN).

The processed data were then converted into Seurat (34, 35) (v5.0.1) for visualization. Local neighborhoods were computed with ‘FindNeighbors’ (dimension = 17), followed by clustering with ‘FindClusters’ (resolution = 0.7). Dimensional reduction was visualized using uniform manifold approximation and projection (UMAP) algorithm. scCustomize (36) (v2.1.2) package in R (v4.3.2) was applied to generate dot plots and feature plots.

### Cell-type identification

An h5ad file was created using the anndata (v0.10.5) package in R and then mapped to a reference 10x Whole Mouse Brain (CCN20230722). The hierarchical mapping algorithm in MapMyCells (RRID:SCR_024672; https://portal.brain-map.org/atlases-and-data/bkp/mapmycells) (37) was chosen to get the broad cell class names, subclass names, and their respective bootstrap probability, which were aligned with the cell barcodes of the Seurat object. As an initial quality control step, cells expressing more than the 99th percentile of the number of genes were removed, and classes with fewer than 10 cells were excluded. Namely, IT (intratelencephalic projecting)- ET (extratelencephalic projecting)-Glut (Glutamatergic neurons), CR (Cajal–Retzius) Glut, NP (near-projecting)-CT (corticothalamic projecting)-L6b Glut, CTX (Cortex)-MGE (medial ganglionic eminence) Gaba (GABAergic neurons), CTX-CGE (caudal ganglionic eminence) Gaba, IMN (immature) Gaba, Astrocytes, OPC (Oligodendrocyte progenitor cells)-Oligo (Oligodendrocytes), OEC (Olfactory ensheathing cells), Vascular and Immune cells. After filtering, cells were manually grouped into the first three main categories—excitatory subtypes, inhibitory subtypes, and non-neuronal cells—based on relevant marker genes and the broad class names provided by the 10x Whole Mouse Brain reference.

For subclustering, dimensionality was determined by identifying the first principal component at which the cumulative explained variance exceeded 90%. Clustering resolution was selected by evaluating resolution-dependent patterns using the Clustree package (38) (v0.5.1). Excitatory cells were subclustered into 8 main groups (15 dimensions, resolution = 0.2): L2/3 IT, L4/5 IT, L5 IT, L6 IT, L5/6 IT, L5 ET, L6 CT, L5 NP, and CR Glut. Inhibitory neurons were subclustered into 10 groups (12 dimensions, resolution = 0.1): Lamp5 Gaba, Sncg Gaba, Vip Gaba, Sst Gaba, Pvalb Gaba, IMN Frmd7 Gaba 1, IMN Frm7 Gaba 2, IMN Thsd7b Gaba, IMN Trdn Gaba, and IMN Dopa-Gaba. Non-neuronal (NN) cells were subclustered to 6 groups (18 dimensions, resolution = 0.1): Microglia NN, Astrocytes NN, Vascular NN, OPC NN, Oligodendrocytes NN and OEC NN.

### Spearman correlation analysis

Normalized expression matrices were obtained for WT and HOM, and *Meis2* expression values were extracted for analysis. Spearman correlation between *Meis2* and all other genes was calculated using the cor() function in R (v4.3.2). Genes with a correlation coefficient ≥ 0.3 were considered significantly correlated and retained for further analysis. Correlation values were visualized using a bar plot generated with ggplot2.

### Predicting RNA velocity using scVelo

RNA velocity was estimated using the scVelo package (v0.3.2) (39). The spliced and unspliced mRNA counts are identified using Velocyto (v0.17.17) (40) with default settings. The first and second moments of spliced and unspliced counts were computed using the scVelo function scvelo.pp.moments with n_neighbors set to 30. The genes that explain the resulting vector field and inferred lineages were identified by scv.tl.rank_velocity_genes.

The velocity vectors were calculated using the scvelo.tl.velocity function, employing the stochastic model of RNA velocity. Velocity confidence intervals were estimated using the scvelo.tl.velocity_confidence function. Velocity embeddings were visualized by projecting the velocity vectors onto UMAP, using the scvelo.pl.velocity_embedding_stream and scvelo.pl.velocity_embedding_grid functions.

### Differential expression analysis and enrichment analysis

To identify genotype-associated differentially expressed genes (DEGs) between each genotype, the scanpy.tl.rank_genes_groups()_function was used to the most variable genes (n = 3000), testing differences in expression between groups of cells. A non-parametric two-sided Wilcoxon rank-sum test was performed at the single-cell level to calculate the log-scale fold change and adjusted p-value for each gene. Only DEGs with an absolute average log fold change >0.5 and an adjusted p-value <0.05 (Benjamini-Hochberg procedure) were considered significant. ClueGO (41) (v2.5.10), one of the modules in Cytoscape (v3.10.2), was used for depicting enriched ontologies for DEGs. Protein-protein interactions between DEGs were illustrated by STRING (v2.2.0) analysis (42) in Cytoscape.

### Cell-cell interaction analysis

CellPhoneDB v5 (43) was used to infer potential ligand-receptor interactions between cell subsets. For each genotype, CellPhoneDB was run separately on the corresponding h5ad file. To focus on biologically relevant interactions, only differentially expressed genes that are significant (Benjamini-Hochberg procedure, FDR < 0.05) were included. These DEGs were used to restrict ligand-receptor pairing, ensuring that predicted interactions reflected genotype-specific transcriptional changes. The predicted interactions were further processed to show the mean of interaction scores, and then the top 5% percentile of interactions based on mean expression level were extracted to plot using the OmicVerse package (v1.6.8) (44).

### RNAscope

After mice were perfused with 4% paraformaldehyde (PFA), brains were removed and post-fixed with 4% PFA overnight. Brains were transferred to 30% sucrose in 1X PBS solution, followed by immersion in O.C.T compound (Sakura Finetek, #4583). 30 μm brain tissue sections were generated on a cryostat. RNAscope was performed following the manufacturer’s protocol for the RNAscope Multiplex Fluorescent Detection Reagents v2 kit (ACD, #323110). The Gad1 (#400951-C2) and Meis2 probes (#436371) were used for co-localization studies. Quantification of RNAscope signals was done using QuPath 0.5.1 (45).

### Statistics

Statistical analyses for each experiment were performed in R (v4.3.3) or GraphPad Prism (v10.1.2). A Shapiro-Wilk normality test was performed to test normal distribution. Two-tailed Student’s t-tests (parametric) or unpaired two-tailed Mann-Whitney U-tests (non-parametric) were used for single comparisons between two groups. For the 3-chamber test, a repeated measures two-way ANOVA test with Tukey HSD correction was performed to compare the two independent variables. All data were expressed as mean ± SEM, with statistical significance determined at p values < 0.05. In detail, p ≥ 0.05 is indicated as ns (no significance), p < 0.05 is indicated as one asterisk (*), p < 0.01 is indicated as two asterisks (**), and p < 0.001 is indicated as three asterisks (***) in all figures.

## Results

### Cell type annotation of single-nucleus transcriptomes in the mPFC of Scn2a WT and HOM mice

To study the impact of *Scn2a* deficiency on the molecular profile of the cells within the mPFC, we collected mPFC samples from WT and *Scn2a* HOM mice and conducted single-nucleus RNA sequencing (snRNA-seq) (**Figure 1A**). We pooled two male and two female mice for each genotype (4 WT and 4 HOM in total) and analyzed transcription from 29,301 nuclei after quality control preprocessing (**Fig. S1A**). For cell-type annotations, we applied MapMyCells (37) to leverage a comprehensive dataset of whole mouse brains from the Allen Brain Institute (**Table S1**). To evaluate the confidence of cell type assignments, we examined the bootstrapping probabilities generated by MapMyCells. This score reflects the proportion of bootstrap iterations in which an annotated cell maps to the same node within the reference cell type taxonomy, providing a measure of classification reliability (37). Most annotated cells exhibited high bootstrapping scores near 1.0 (**Fig. S1B-D**), indicating consistent and confident mapping across hierarchical levels of the taxonomy.

**Figure 1.**
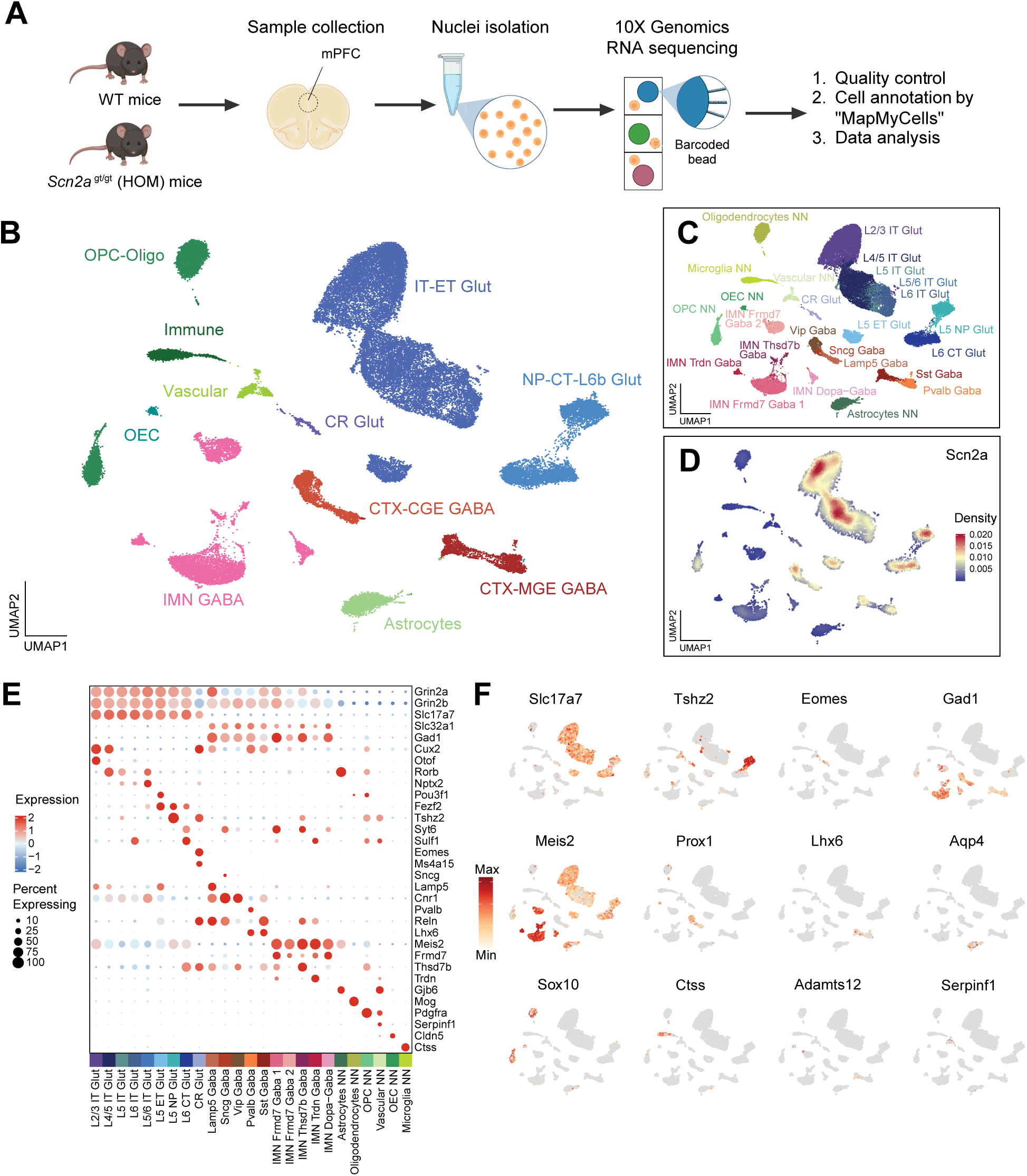
MapMyCells analysis identified 11 classes with 25 subclasses in single nucleus RNA-sequencing in the mPFC of *Scn2a*-deficient mice (A) Schematic diagram for experimental pipelines created with BioRender.com. Medial prefrontal cortex (mPFC) tissue was collected from WT and homozygous *Scn2a*-deficient (HOM) mice for single-nucleus RNA sequencing. Downstream analysis included quality control, cell type annotation using MapMyCells (RRID:SCR_024672), and transcriptomic analysis. (B) Comprehensive uniform manifold approximation and projection (UMAP) demonstrating class annotations based on MapMyCells, including glutamatergic neurons (IT-ET Glut, NP-CT-L6b Glut, CR Glut), GABAergic neurons (CTX-CGE GABA, CTX-MGE GABA, IMN GABA), astrocytes, oligodendrocyte lineage (OPC-Oligo), olfactory ensheathing cells (OEC), immune cells, and vascular cells. (C) UMAP embedding visualizing fine subclass annotations based on MapMyCells. (D) Density plot showing regional density distribution of nuclei across subclasses for *Scn2a* gene. Note that *Scn2a* gene is expressed in glutamatergic neurons and GABAergic neurons. (E) Dot plot depicting expression patterns of canonical marker genes across annotated subclasses. *Grin2a, Grin2b, Slc16a7, Cux2, Otof, Rorb, Nptx2, Pou3f1, Fezf2, Tshz2, Syt6, Sulf1* for glutamatergic neurons; *Slc32a1, Gad1, Eomes, Ms4a15, Sncg, Lamp5, Cnr1, Pvalb, Reln, Lhx6, Meis2, Frmd7, Thsd7b, Trdn* for GABAergic neurons; *Gjb6, Mog, Pdgfra, Serpinf1, Ctss, Cldn5* for non-neuronal cells. The color gradient from blue (low) to red (high) represents scaled expression, and dot size indicates the percentage of cells expressing each gene. (F) Feature plots of selected marker genes overlaid on the UMAP, illustrating their distribution across specific neuronal and non-neuronal populations. The color gradient from light orange (low) to dark red (high) represents relative expression levels, while gray dots represent cells with no expression.

Uniform manifold approximation and projection (UMAP) dimensionality reduction and Leiden clustering of the transcriptomic datasets identified 11 distinct cell classes and 25 subclasses (**Figure 1B-C**). We observed that *Scn2a* is primarily expressed in intratelencephalic projecting (IT)-extratelencephalic projecting (ET) and near-projecting (NP)-corticothalamic projecting (CT)-L6b glutamatergic neurons, as well as in CTX- caudal ganglionic eminence (CGE) GABA neurons and CTX-medial ganglionic eminence (MGE) GABA neurons, consistent with known expression patterns of the *Scn2a* gene (46, 47) (**Figure 1D**).

We profiled the expression of established marker genes for major mPFC cell types: *Grin2a*, *Grin2b*, and *Slc17a7* for glutamatergic neurons; *Slc32a1* and *Gad1* for GABAergic neurons; *Cux2, Otof, Rorb, Nptx2*, and *Pou3f1* for IT-ET Glut neurons; *Fezf2, Tshz2, Syt6*, and *Sulf1* for NP-CT-L6b Glut neurons; and *Eomes* and *Ms4a15* for Cajal– Retzius (CR) Glut neurons. *Sncg, Lamp5,* and *Cnr1* were specifically expressed in CTX-CGE GABA neurons, while *Pvalb* and *Lhx6* were expressed in CTX-MGE GABA neurons (**Figure 1E**). Notably, we observed a large subpopulation of *Meis2*+ immature GABAergic neurons identified by co-expression of *Meis2, Frmd7, Thsd7b,* and *Trdn,* along with *Gad1*, a gene responsible for synthesizing GABA. In addition to neurons, we annotated non-neuronal (NN) cells based on marker gene expression: *Gjb6* for astrocytes, *Mog* for oligodendrocytes, *Pdgfra* for oligodendrocyte progenitor cells (OPCs), *Serpinf1* for vascular cells, *Cldn5* for olfactory ensheathing cells (OECs), and *Ctss* for microglia (**Figure 1E**). Cell type annotations aligned with canonical markers, as shown in feature plots of gene expression across cell types for glutamatergic, GABAergic neurons, and non-neuronal cell types (**Figure 1F**). Together, these results demonstrate the identification of distinct neuronal and non-neuronal subtypes in the mPFC using MapMyCells (RRID:SCR_024672) (37). A complete list of class and subclass annotations for cell types is provided in **Table S2**.

### Meis2+ Immature GABAergic neurons are increased in the mPFC of Scn2a-deficient mice

As we observed *Meis2*+ immature GABAergic neurons with expression of *Thsd7b, Trdn*, and *Frmd7* in our dataset, we sought to verify the expression of these marker genes using reference datasets. The Allen Brain Cell Atlas (https://knowledge.brain-map.org/abcatlas), which provides the single-cell spatial transcriptome of the adult mouse brain (37), confirmed high expression of these genes in OB-IMN-GABA neurons (**Fig. S2A**), supporting that the population we identified corresponds to this subtype. OB-IMN-GABA neurons originate from the lateral ganglionic eminence (LGE) and migrate to the olfactory bulb (OB) (26, 48). Although these neurons are primarily localized in the anterior olfactory nucleus (AON), small populations are also found in the striatum, lateral septal complex, and prelimbic/infralimbic regions of the mPFC (**Fig. S2B**) (49). Consistent with these reference data, *Meis2*+ immature GABAergic neurons have previously been reported in the mPFC of adult WT mice using snRNA-seq (37, 49).

Interestingly, UMAP in our dataset revealed an increased IMN GABA neuron population in the *Scn2a^gt/gt^* (HOM) mice mPFC compared to WT (**Figure 2A**). A summary bar plot indicated that the proportion of IMN GABA neurons in mPFC was 2.99% in the WT but increased to 19.39% in the HOM, as highlighted in red (**Figure 2B**). Moreover, *Meis2* is primarily expressed in the IMN GABA class and strongly co-expressed with the canonical GABAergic marker *Gad1* in our dataset (**Figure 1E**). A comparison between genotypes showed that the number of *Meis2*+*Gad1*+ cells was increased in HOM mice relative to WT. (**Fig. S2C**), confirming the expansion of this immature inhibitory neuron population in the *Scn2a*-deficient mice.

**Figure 2.**
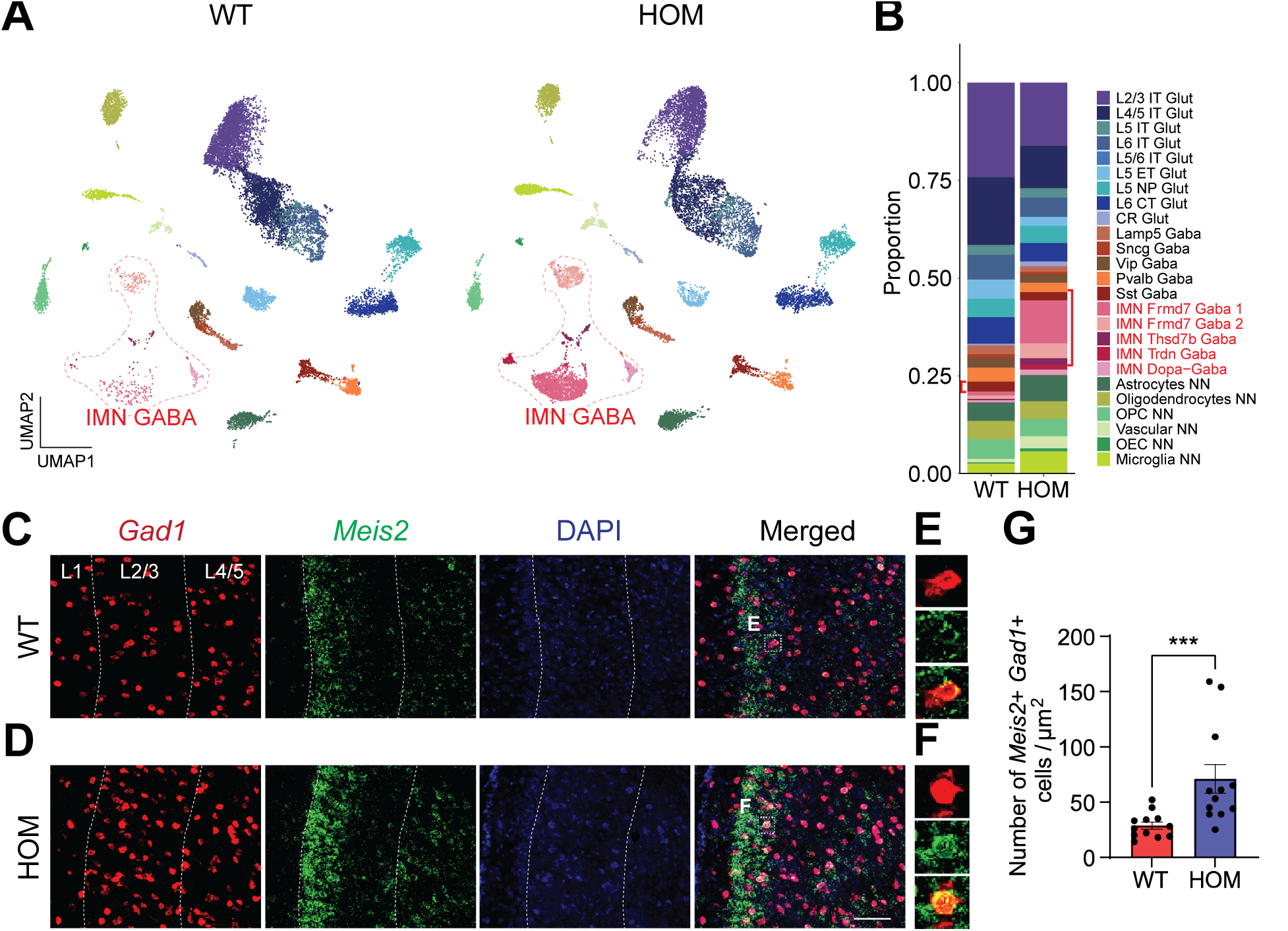
The population of *Meis2*-expressing GABAergic neurons is increased in *Scn2a*- deficient mice (A) UMAP for WT and HOM showing differential cell population. Note that the population of IMN GABA neurons in red areas was increased in HOM. (B) Stacked bar plot representing cell-type-proportions in WT and HOM. IMN GABA neuron cell proportion is highlighted in red. (C, D) Representative images of RNAScope results in WT (C) and HOM (D). Red, *Gad1*; Green, *Meis2*; Blue, DAPI. Layers 1, 2/3, 4/5 of the mPFC are labeled. Scale bar = 100 µm (E, F). Magnified images of cells expressing both *Meis2* (green) and *Gad1* (red) gene in WT (E) and HOM (F). (G) *Meis2* and *Gad1* co-expressing cells were increased in HOM compared to WT. p < 0.001, ***. Mann-Whitney test with Shapiro-Wilk test. WT (n = 12 from 4 animals) and HOM (n = 12 from 4 animals).

To validate the transcriptomic finding of increased *Meis2*+ IMN GABA neuron proportion in HOM mice, RNAScope *in situ* hybridization was performed using *Meis2* and *Gad1* probes. The RNAScope results in the mPFC region indicated that the *Meis2*+ IMN GABA neuron population increased in adult HOM mice compared to WT (**Figure 2C, 2D**). Co- localization of *Gad1*+ and *Meis2*+ signals in each cell were also assessed (**Figure 2E, 2F**). The number of cells expressing both *Meis2* and *Gad1* in the region of interest was higher in HOM compared to WT (mean = 28.8 cells/μm^2^ for WT, mean = 71.1 cells/μm^2^ for HOM, p = 0.0010, Mann-Whitney U test) (**Figure 2G**). The increased number of *Meis2*+ and *Gad1*+ IMN GABA neurons observed in the adult stage of HOM mice, along with the transcriptomic data, collectively indicates that *Scn2a* deficiency leads to changes in GABAergic neuron populations.

To further investigate transcriptional networks associated with *Meis2* in the mPFC, we performed Spearman correlation analysis to identify genes whose expression patterns were significantly correlated with *Meis2.* Overall, more genes showed positive (red bars) than negative (blue bars) correlations with *Meis2*, indicating that most *Meis2*-associated transcriptional relationships were positively coupled (**Fig. S3A-B**). Notably, a larger number of strongly correlated genes was observed in HOM mice compared to WT, suggesting that *Scn2a* deficiency may enhance transcriptional coupling or co-regulation with *Meis2*. Many of these genes in HOM are involved in synaptic function, such as *Dlgap1, Synpr*, and *Gria2*, as well as ion channel regulation, including *Kcnj3*, *Kcnd2*, and *Gabrb3*. Several transcription factors, including *Pbx1, Hdac9*, and *Pbx3*, also showed stronger correlations with *Meis2* in HOM compared to WT. These findings suggest that *Scn2a* deficiency may alter the transcriptional context of *Meis2*, contributing to broader dysregulation of synaptic and ion channel-related genes.

### Genes altered in the transcriptomic profile of Scn2a-deficient mice are associated with neuronal development

A recent study reported that loss of *Scn2a* can affect neurogenesis and neuronal differentiation (50). Therefore, we hypothesized that *Scn2a* deficiency induced transcriptional changes during development, leading to aberrant cell fate decisions and increased differentiation of immature GABAergic neurons in adulthood. To test this hypothesis, we employed scVelo, a computational method that models gene expression dynamics using snRNA-seq data (39). Specifically, scVelo estimates RNA velocity, a vector representing the direction and speed of a cell’s transcriptional state, by comparing levels of unspliced and spliced RNA transcripts (39, 51). These velocity estimates are then used to infer latent time from pseudotemporal ordering, which places each cell along a developmental trajectory according to its transcriptional maturity (40, 52).

Integration of RNA velocity with UMAP embeddings revealed overall comparable developmental trajectories from immature (IMN) to mature (CTX-MGE and CTX-CGE) GABAergic neuron subtypes between WT and HOM (**Fig. S4A**). Despite overall similarities in inferred lineage flow between WT and HOM mice, the distribution of cell states differed across the trajectory. IMN GABAergic neurons were markedly increased in HOM mice and were positioned at earlier stages of the inferred trajectory (**Fig. S4B**). Notably, IMN GABA neurons and CTX-MGE/CGE neurons formed distinct clusters on the latent time map, supporting clear separation between immature and mature GABAergic populations. Together, these observations suggest that *Scn2a* deficiency may alter the distribution of GABAergic cell states, leading to an expansion of immature neurons and a possible compensatory change in more mature subtypes.

Next, we investigated putative driving genes, which exert a strong influence on the direction and rate of cell state transitions along dynamic differentiation trajectories. We plotted the top-ranked driving genes based on their differential RNA velocity dynamics between HOM and WT samples (**Table S3**). To examine the molecular function of driving genes, we focused on those that were preferentially expressed in IMN GABA neurons. Given their potential involvement in neurodevelopment, we hypothesized that these genes may regulate key neurodevelopmental processes, including neuronal proliferation, differentiation, and patterning. Notably, several neuronal development-related genes such as *Ano4, Pbx1, Ptpro*, and *Cnln5* were upregulated in IMN GABA neurons in HOM compared to WT (**Fig. S4C**). This upregulation in HOM suggests that *Scn2a* deficiency may delay or misdirect transcriptional programs required for proper neuronal maturation and regional specification, contributing to a shift in GABAergic cell fate.

To validate the developmental stage of GABAergic neurons, we assessed the ratio of *Nkcc1* (sodium potassium chloride cotransporter 1) and *Kcc2* (potassium chloride co- transporter 2) gene expression. This *Nkcc1*/*Kcc2* ratio is critical for regulating neuronal chloride homeostasis and modulating GABAergic neurotransmission (53, 54) and is commonly used as an indicator of GABAergic neuron maturation (55, 56). As GABAergic neurons mature, *Nkcc1/Kcc2* expression gradually reduces (57, 58). HOM mice showed an increased *Nkcc1*/*Kcc2* ratio compared to the WT only in IMN GABA neurons, but not in other mature GABAergic populations (**Fig. S4D**). This finding further supports that IMN GABA neurons remain in an immature state in HOM. Together, these findings provided insights into changes in transcriptomic dynamics of aberrantly increased immature GABAergic neurons in *Scn2a*-deficient mice.

### Scn2a-deficient GABAergic neurons showed disruption in postsynaptic membrane and neurotransmitter activity regulation

Our scVelo analysis demonstrated that an immature neuronal population in the inhibitory lineage is shaped by distinct transcriptional programs inferred from latent time and driving gene dynamics. To investigate cell type-specific transcriptional changes associated with *Scn2a* deficiency, we performed differential expression analysis across distinct cell classes. Significantly differentially expressed genes (DEGs) were identified using thresholds of false discovery rate (FDR) < 0.05 and |log fold change (logFC)| > 0.5. We then quantified the number of DEGs in each class (**Figure 3A, Table S4**). This class-level pseudo-bulk approach, aggregating individual nuclei into cell type groups and performing pairwise analyses between WT and HOM genotypes, enabled us to identify genes and pathways disrupted in the major neuronal subtypes within the mPFC. Among all classes, IT-ET Glut neurons showed the highest number of DEGs (**Figure 3A**), indicating a pronounced transcriptional impact of *Scn2a* deficiency in this population, possibly related to *Scn2a*’s high expression in glutamatergic neurons.

**Figure 3.**
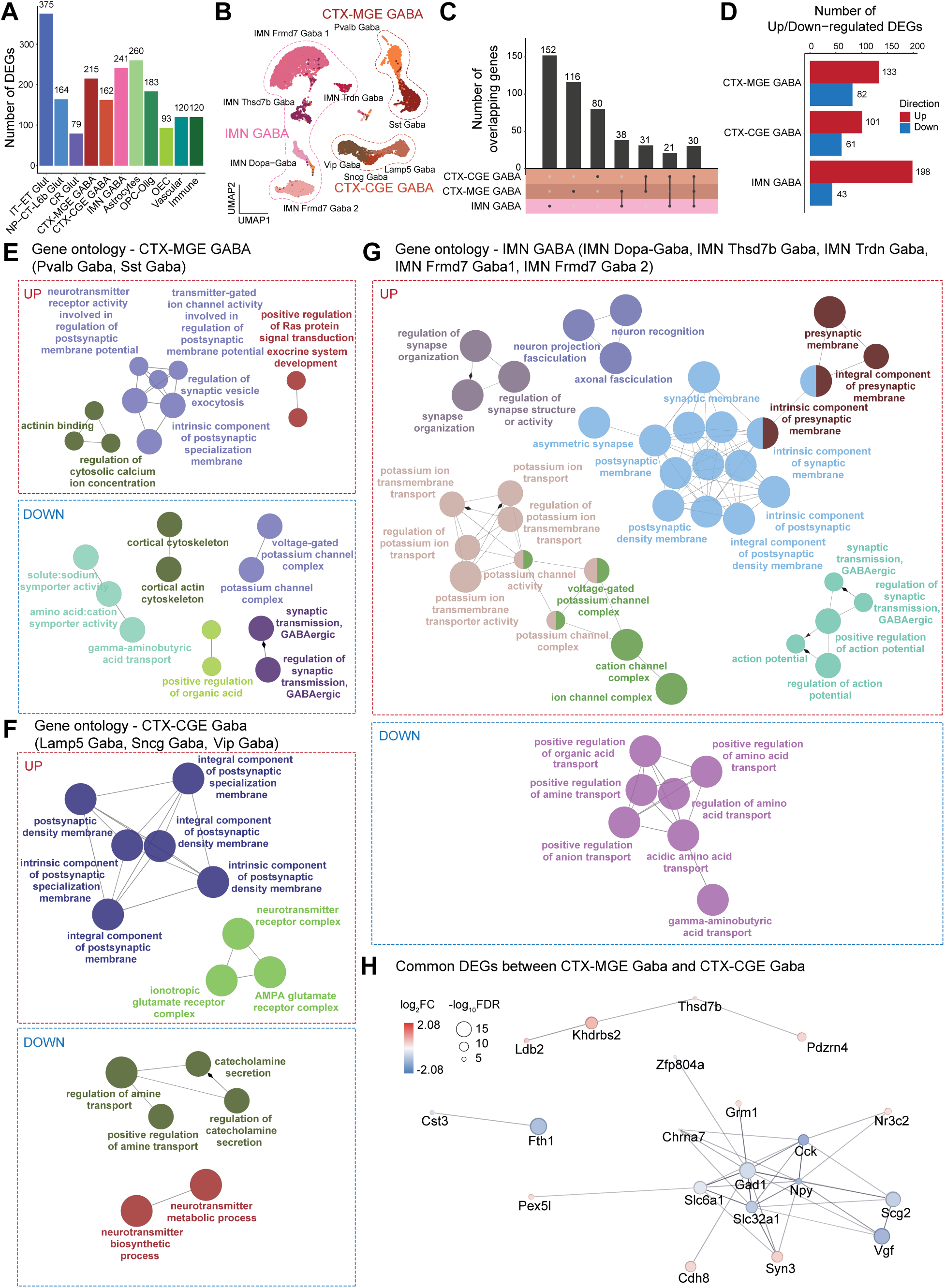
Differentially expressed gene (DEG) analysis showed a disruption in ion channels and neurotransmitter activity in *Scn2a*-deficient GABAergic neurons (A) Bar plot representing the number of significant DEGs in each class type. (B) UMAP of subset GABAergic neurons. (C) UpSet plot showing the number of common and unique genes shared across GABAergic neuron classes between HOM and WT. Rows in the matrix represent the class types, and columns indicate the intersections among these class types. (D) Number of up/down- regulated DEGs across Gaba neuron classes. Gene ontology (GO) enrichment analyses of upregulated and downregulated DEGs from: (E) CTX-MGE GABA neurons, (F) CTX-CGE GABA neurons, and (G) IMN GABA neurons. (H) Network visualization of shared DEGs between CTX-MGE GABA and CTX-CGE GABA neurons analyzed by STRING. Node size corresponds to log10(FDR), and color indicates log2 fold change direction. Red, upregulated; Blue, downregulated.

To investigate transcriptomic changes within GABAergic neuron subtypes, we subclustered GABA neurons into three major classes: CTX-MGE GABA, CTX-CGE GABA, and IMN GABA. (**Figure 3B**). To account for differences in IMN GABA cell numbers between genotypes, DE analysis was performed using normalized gene expression per cell. Like the increase in the IMN GABA population in HOM mice compared to WT (**Figure 2**), the IMN GABA class exhibited the greatest number of significant DEGs among GABAergic subclasses, followed by CTX-MGE GABA and CTX-CGE GABA neurons (**Figure 3C, 3D**). Notably, 30 DEGs were shared across all three GABAergic classes, including *Pex5l, Nr3c2, Syn3*, and *Slc6a1,* which are involved in synapse regulation. These results suggest that *Scn2a* deficiency may induce both subtype-specific and shared transcriptomic alterations across GABAergic neuron classes.

To uncover the functional implications of DEGs in GABAergic neurons identified above, we performed ClueGO functional enrichment analysis (41) in Cytoscape and identified ontologies related to brain function and neuronal regulation. The ClueGO module visualized enriched gene ontology (GO) terms across biological process (BP), cellular component (CC), and molecular function (MF) networks for DEGs identified in IMN GABA, CTX-MGE GABA, and CTX-CGE GABA (**Table S5-6**).

In the CTX-MGE GABA class, 133 upregulated DEGs were enriched in GO terms related to the regulation of postsynaptic membrane potential (**Figure 3E**). 82 downregulated DEGs were associated with disrupted processes such as regulation of GABAergic synaptic transmission and voltage-gated potassium channel complex (**Figure 3E**). Similarly, in the CTX-CGE GABA class, 101 upregulated DEGs were enriched in postsynaptic density and membrane specialization, while 61 downregulated DEGs were linked to regulation of amine transport and neurotransmitter biosynthetic process (**Figure 3F**). For IMN GABA class, 198 upregulated DEGs formed large, enriched networks involved in the regulation of postsynaptic density membrane, action potential, and potassium ion transmembrane transport (**Figure 3G**). 19 downregulated DEGs were linked to the regulation of amino acid transport (**Figure 3G**).

To identify genes most impacted by *Scn2a* deficiency across mature GABAergic subtypes, we next focused on commonly dysregulated genes between the CTX-MGE and CTX-CGE GABA classes. Among the 31 shared DEGs, STRING analysis (42) revealed downregulation of GABA transport-related genes including *Gad1* and *Slc32a1*, and upregulation of synaptic genes including *Cdh8* and *Syn3* in HOM (**Figure 3H, Table S7**). Given that *Scn2a* is expressed in both CTX-MGE GABA and CTX-CGE GABA neurons (**Figure 1B and D**), these results suggest that *Scn2a* deficiency might lead to compensatory upregulation of synaptic genes in mature GABA neurons, resulting in disruption of postsynaptic density regulation and membrane specialization.

### Scn2a-deficient glutamatergic neurons have differential gene expression of transcripts involved in synapse transmission and ion transport

Since *Scn2a* is primarily expressed in excitatory neurons (**Figure 1B and D**) (2, 14), we next focused on transcriptomic changes in excitatory neurons, whose population sizes remain comparable between WT and HOM. Specifically, we examined whether their gene expression differences mirrored the patterns observed in inhibitory neurons.

We subclustered the excitatory neuron population in the UMAP and focused on cell types within the IT-ET Glut and NP-CT-L6b Glut class (**Figure 4A**). The IT-ET Glut class showed the highest number of DEGs among glutamatergic subtypes in our analysis (**Figure 4B, 4C**). GO enrichment analysis revealed that 205 DEGs upregulated in IT-ET Glut neurons of HOM mice were significantly enriched in action potential regulation and plasma membrane cell adhesion molecules, while 170 downregulated DEGs were associated with amino acid transport and postsynaptic organization (**Figure 4D, Table S5-6**). For the NP-CT-L6b Glut class, 110 upregulated DEGs were enriched in nuclear receptor activity, while 54 downregulated genes were linked to glutamate receptor activity (**Figure 4E**).

**Figure 4.**
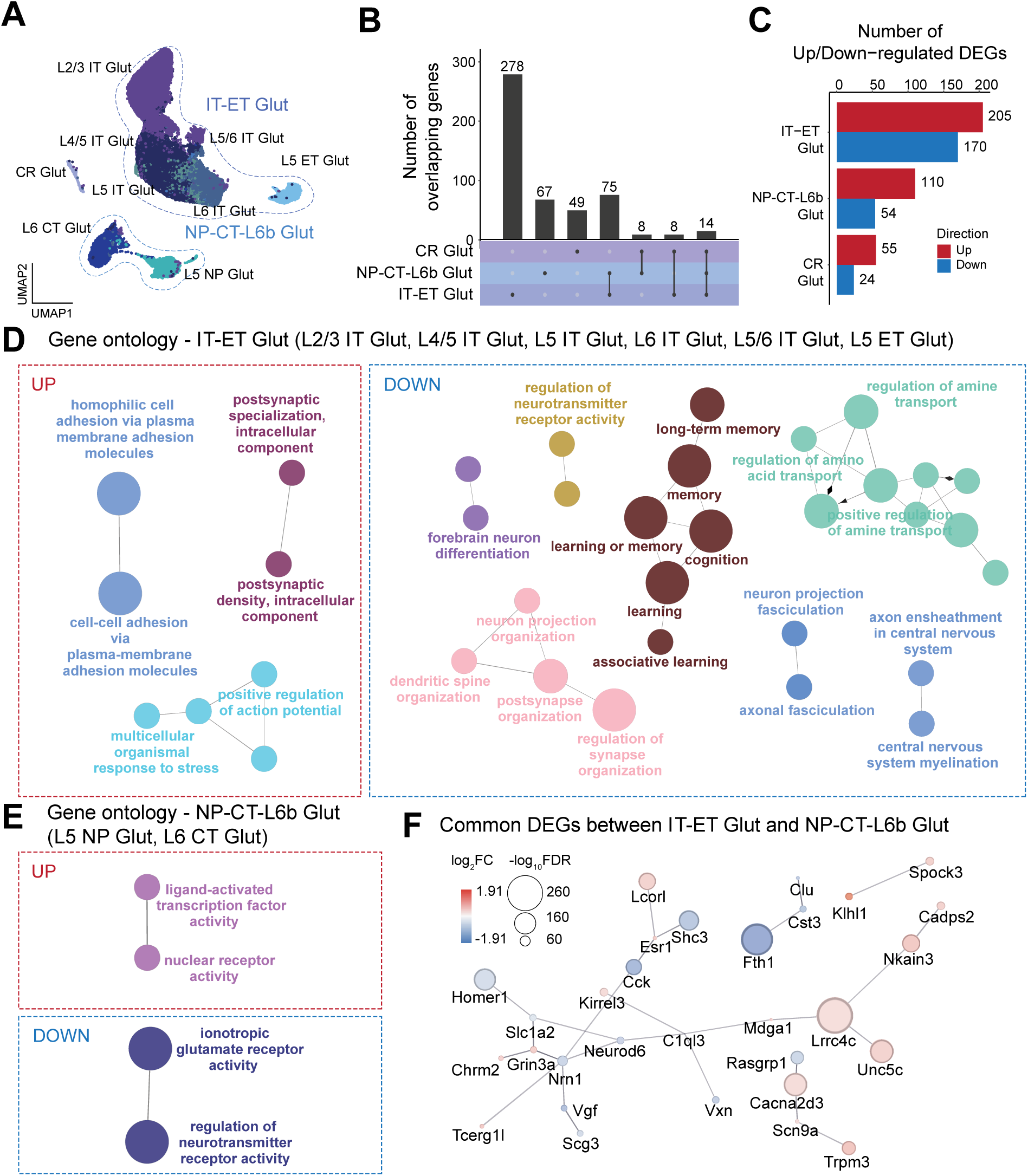
Differentially expressed genes in *Scn2a*-deficient glutamatergic neurons are enriched in synaptic organization and neuronal regulation ontologies (A) UMAP of subset glutamatergic neurons. (B) UpSet plot showing the number of common and unique DEGs in Glut neurons between *Scn2a* HOM and WT. Rows in the matrix represent the class types, and columns indicate the intersections among these class types. (C) Number of up/down-regulated DEGs by Glut neuron types. Enriched gene ontology (GO) of upregulated and downregulated unique DEGs from: (D) IT-ET Glut neurons, and (E) NP-CT-L6b Glut neurons. (F) Protein-protein interaction network of DEGs commonly expressed between IT-ET Glut neurons and NP-CT-L6b Glut neurons analyzed by STRING (v2.2.0) in Cytoscape.

To identify common molecular changes across excitatory neuron subtypes, we performed STRING network analysis on DEGs shared between the IT-ET Glut and NP-CT-L6b Glut classes. This analysis revealed a connected gene network of hub genes implicated in neuronal development, synaptic signaling, and glutamate transmission, including *Neurod6, Nrn1, Grin3, Slc1a2*, and *Lrrc4c* (**Figure 4F, Table S7**). These findings suggest a coordinated dysregulation of core functional pathways in excitatory neurons of HOM mice. The presence of shared DEGs across distinct excitatory classes indicates that *Scn2a* deficiency induces convergent transcriptional changes that may broadly impair glutamatergic signaling in both L2/3 and L4/5 cortical layers. Together with DE analysis, these results suggest that *Scn2a* is essential for maintaining excitatory neuron function and further suggest that its loss might disrupt key regulatory networks necessary for proper cortical circuit activity.

### Glial cells show differential gene expression of transcripts associated with forebrain neuron development and synapse regulation in response to Scn2a deficiency

We identified DEGs and enriched pathways in non-neuronal cells to assess the transcriptional impact of *Scn2a* deficiency. UMAP subclustering revealed distinct class types corresponding to Astrocytes, OPC-Oligo (oligodendrocyte progenitor cells and oligodendrocytes), Immune (microglia), OEC (olfactory ensheathing cells), and Vascular classes (**Figure 5A**). By focusing on the three major glial types, we identified overlapping and unique DEGs across Astrocytes, OPC-Oligo, and Immune classes. Astrocytes class exhibited the largest number of unique DEGs, followed by Olig-OPC and Immune classes. (**Figure 5B, 5C**).

**Figure 5.**
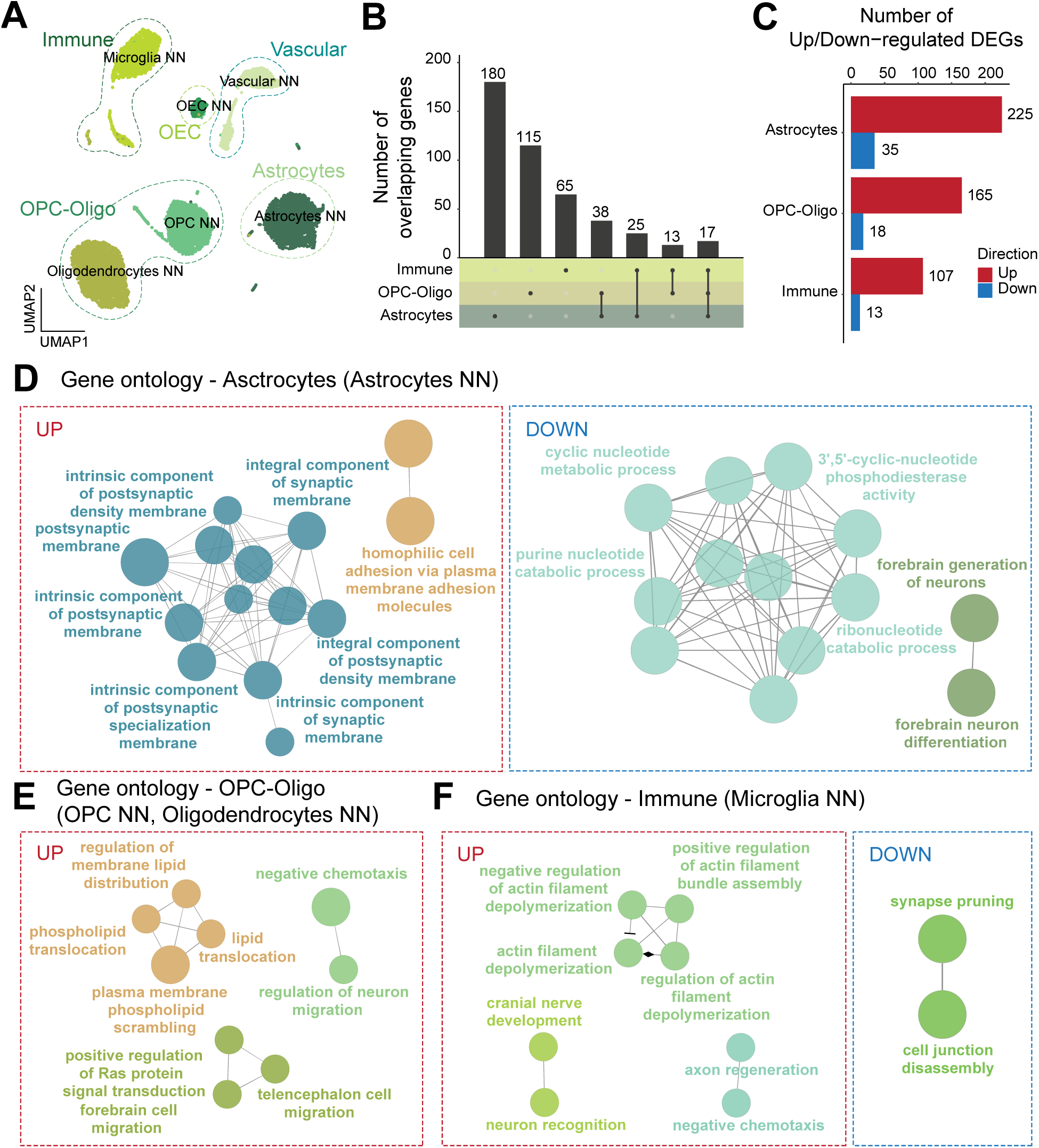
Differentially expressed genes in *Scn2a*-deficient glial cells are enriched in ligand-gated ion channel activity and membrane regulation ontologies (A) UMAP of subset non-neuronal cells. (B) UpSet plot showing the number of common and unique DEGs in three major types (Astrocytes, OPC-Oligo, Immune) of glial cells between *Scn2a* HOM and WT. Rows in the matrix represent the class types, and columns indicate the intersections among these class types. (C) Number of up/down-regulated DEGs by glial cell types. Enriched gene ontology (GO) of upregulated and downregulated unique DEGs from: (D) Astrocytes, and (E) OPC-Oligo. (F) Enriched GO of upregulated and downregulated unique DEGs from Immune.

GO enrichment analysis was then performed for upregulated and downregulated DEGs within each glial class. Among them, Astrocytes showed the most extensive enrichment networks compared to OPC-Oligo and Immune cells (**Figure 5D**). In Astrocytes, upregulated genes were associated with postsynaptic density membrane, suggesting potential alterations in neuron-glia synaptic interactions. Downregulated genes were enriched in pathways related to forebrain neuron differentiation, likely reflecting a disruption in astrocyte-mediated support for neuronal development or maintenance (**Figure 5D**). In OPC-Oligo, upregulated DEGs showed enrichment in membrane lipid distribution and neuron migration, while no significant enrichment was observed in downregulated genes (**Figure 5E**). For the Immune class, upregulated DEGs were associated with the regulation of actin depolymerization and nerve development, whereas downregulated DEGs were enriched in synapse pruning (**Figure 5F**). Given that *Scn2a* has very limited expression in glial populations, these transcriptomic changes likely reflect secondary, non-cell autonomous effects in response to *Scn2a* deficiency. Notably, astrocytes appeared to be the most transcriptionally affected glial cell type, highlighting their potential role in modulating neuronal function following *Scn2a* loss.

### Candidate genes in neuronal modulation were restored by FlpO genetic rescue

From our analysis, we identified cell type-specific transcriptomic alterations in glutamatergic and GABAergic neurons caused by *Scn2a* deficiency and wanted to explore whether re-expression of *Scn2a* in adulthood could reverse these transcriptional changes. We performed a genetic rescue experiment by systemically administering AAV-PHP.eB-FlpO via tail vein injection, which excises the gene trap cassette and restores *Scn2a* expression (17). To evaluate the functional impact of gene restoration, we performed a three-chamber sociability test before tissue collection (**Figure 6A**). Consistent with previous findings, *Scn2a* HOM mice exhibited impaired social interaction, spending less time with a social target than WT controls (30). Genetic restoration via FlpO in HOM-FlpO mice rescued this deficit, restoring normal sociability (**Figure 6B**). As expected, transcriptomic analyses in HOM-FlpO mice revealed that adult *Scn2a* restoration did not reduce the expanded IMN GABA population (**Figure 6C**) nor their earlier latent time characteristics (data not shown), since developmental alterations in cell fate and cellular composition are difficult to reverse once established. Therefore, the observed rescue of social behavior may be attributed to transcriptomic changes within established cell populations.

**Figure 6.**
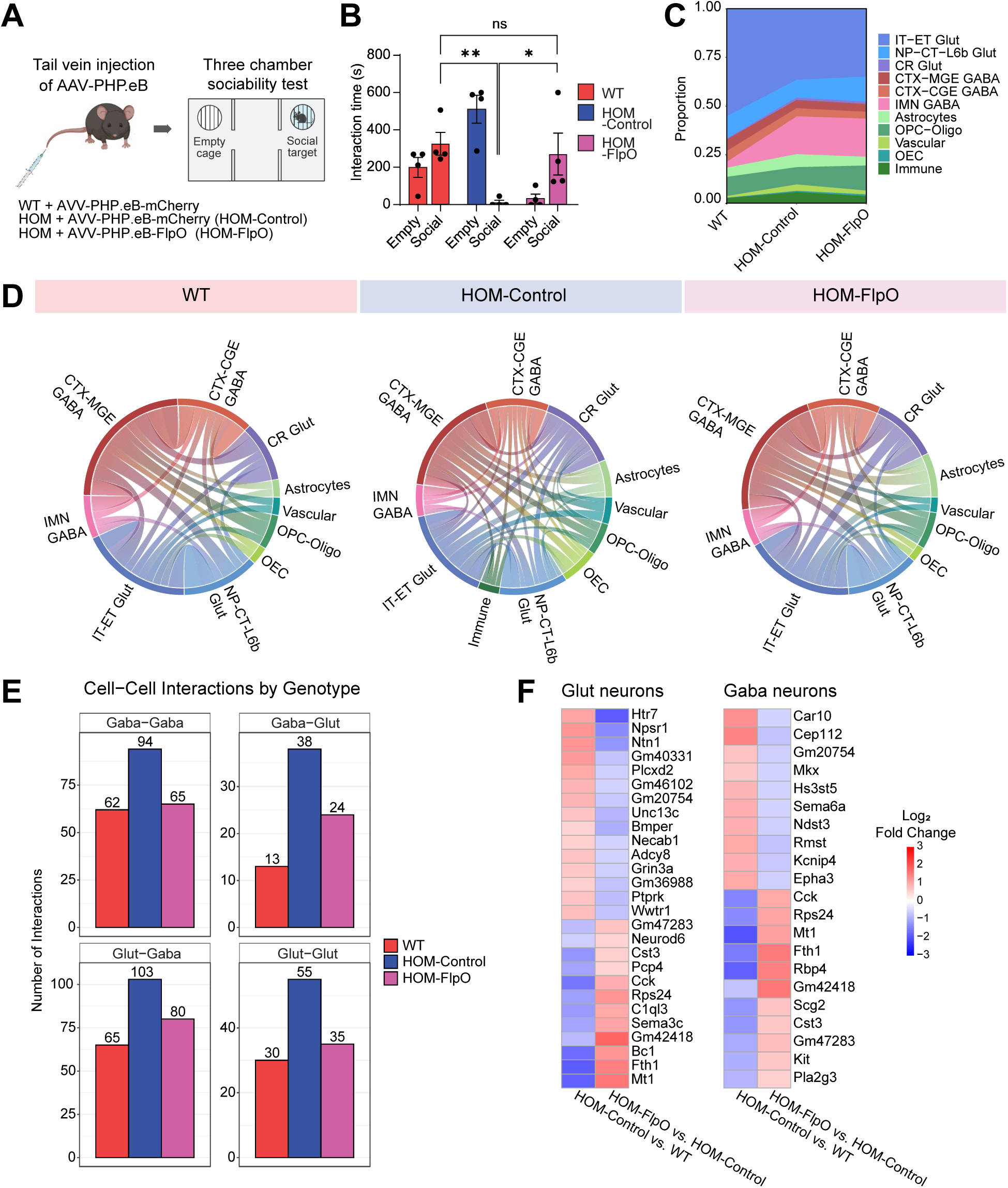
Restoration of *Scn2a* expression improves social behavior and neural communication networks in *Scn2a*-deficient mice. (A) Schematic diagram of experimental design for tail vein injection of AAV-PHP.eB-mCherry/FlpO and three-chamber sociability test, created with BioRender.com. (B) Time spent interacting with a social target vs. empty chamber for WT, HOM-Control, and HOM-Flp mice. HOM-Control mice showed social deficit behavior, whereas HOM-FlpO mice showed normalized social behavior. p < 0.05, *; p < 0.01, **; ns, no significance. Two-way ANOVA. (C) Stacked area plot showing the proportions of neuronal and non-neuronal cell classes across genotypes (WT, HOM-Control, HOM-FlpO), color-coded by cell type. The IMN GABA population persists in both HOM and HOM- FlpO groups. (D) Circos plots representing predicted cell-cell communication networks in WT, HOM-Control, and HOM-FlpO. (E) Bar plot displaying the total number of interactions in WT, HOM-Control, and HOM-FlpO between Gaba and Glut neuron pairs (Gaba–Gaba, Gaba–Glut, Glut–Gaba, Glut–Glut). (F) Heatmap of DEGs between HOM-FlpO and HOM groups, showing partial transcriptional normalization in HOM-Control vs. WT and HOM-FlpO vs. HOM-Control. Red, upregulated; Blue, downregulated.

To explore the underlying mechanisms, we first investigated whether altered intercellular communication was modulated by *Scn2a* restoration. Using CellPhoneDB, a tool that utilizes ligand and receptor gene expression databases to predict cell-cell communication (43), we assessed predicted signaling interactions across all neuronal cell types. Intriguingly, HOM mice showed an increased number of predicted interactions compared to WT, which was reduced in HOM-FlpO mice (**Figure 6D, Table S8**). To further dissect cell type-specific communication, we analyzed the number of interactions categorized by source and target neuronal subtypes. *Scn2a* restoration in HOM mice broadly reduced interactions across all GABAergic and glutamatergic cell pairings (GABA-GABA, GABA-Glut, Glut-GABA, and Glut-Glut) (**Figure 6E**). This finding indicates that social rescue may be driven not by reversal of GABAergic neuron composition but rather by normalization of existing aberrant cell-cell communication networks.

Finally, we conducted differential expression analysis in HOM-FlpO vs. HOM to uncover gene expression alteration driven by adult *Scn2a* restoration. We focused on DEGs showing opposite expression changes in WT vs. HOM and HOM vs. HOM-FlpO within glutamatergic and GABAergic neurons. This analysis identified 27 ‘rescued genes’ in glutamatergic neurons and 21 in GABAergic neurons (**Figure 6F, Table S9**). Notably, genes such as *Htr7*, *Grin3a*, and *Neurod6* in glutamatergic neurons, and *Sema6a*, *Scg2*, and *Kcnip4* in GABAergic neurons are linked to synaptic signaling, cell adhesion, and ion channel function. These findings suggest that *Scn2a* gene restoration in adulthood can reverse disease-associated transcriptomic changes by modulating key regulators of neuronal signaling and connectivity without altering developmentally established neuronal composition.

## Discussion

In this study, we employed single-nucleus RNA sequencing to define the cell-type-specific transcriptional landscape of the medial prefrontal cortex (mPFC) in a mouse model of *Scn2a* deficiency. We profiled transcriptomic changes using *Scn2a* homozygous gene trap mice (HOM) and wild-type (WT) controls. By adopting MapMyCells from the Allen Brain Institute (37), cell identities were annotated to reveal major classes, including IT-ET Glut, NP-CT-L6b Glut, CTX-MGE GABA, CTX-CGE GABA, IMN GABA, and non-neuronal cells. Notably, we observed a marked increase in the proportion of IMN GABA neurons in the mPFC of *Scn2a* HOM mice. These cells expressed neuronal development-related genes and showed an elevated *Nkcc1*/*Kcc2* expression ratio, consistent with an immature phenotype. Differential gene expression analysis showed that *Scn2a* deficiency alters genes related to GABAergic synaptic transmission and voltage-gated potassium channels in GABAergic neurons, while glutamatergic neurons are enriched for pathways involving membrane potential and glutamate receptor signaling. These transcriptomic alterations in different cell types may contribute to the cortical hyperexcitability and social deficits observed in the *Scn2a*-deficient mouse model (30).

Most notably, the increased population of IMN GABA neurons is marked by canonical genes associated with early GABAergic neuron development: *Meis2, Trdn, Frmd7,* and *Thsd7b* (*49*) (**Figure 1E**). *Meis2* is a transcription factor involved in neurodevelopment and cell fate specification (59, 60). *Trdn* and *Frmd7*, are also associated with neuronal development: *Trdn* encodes a protein involved in calcium signaling (61), while *Frmd7* is important for axon guidance and neuronal migration (62, 63). *Thsd7b* encodes a membrane component that plays a critical role in actin cytoskeleton reorganization and is involved in post-translational modifications and glycosylation of proteins (64). *Meis2* is a key transcription factor involved in GABAergic neuron differentiation (65), and there are more *Meis2*-correlated genes in *Scn2a*-deficient HOM mice compared to WT (**Fig. S2B)**. Genes highly correlated with *Meis2* were found to regulate synaptic function and ion channel regulation. These findings suggest that elevated *Meis2* expression may contribute to altered neuronal differentiation in the mPFC of *Scn2a*-deficient mice.

Interestingly, a recent single-cell RNA sequencing (scRNA-seq) study of human cortical organoids carrying a nonsense mutation *SCN2A^+/G1744X^* and complete loss-of-function *SCN2A^-/-^* reported the greatest number of differentially expressed genes (DEGs) in excitatory neurons, which aligns with our findings (50). This study further identified an increased proportion of CGE-like inhibitory neurons, confirming cell type–specific impact of *SCN2A* loss. Together with our results, these findings underscore the central role of *Scn2a* in guiding neurogenesis and suggest that interventions restoring *Scn2a* function may have the potential to normalize transcriptional programs.

We observed increased predicted cell-cell communication between identified class types in the mPFC of *Scn2a* HOM mice (**Figure 6**). This finding aligned with increased neural communication networks observed in scRNA-seq data from individuals with autism spectrum disorder (ASD) (66, 67). Functional MRI studies in ASD patients also reported brain hyperconnectivity, which correlates with social deficit severity (68, 69). In this context, our observation that FlpO-mediated *Scn2a* gene restoration in HOM-FlpO mice partially normalizes predicted intercellular communication suggests a possible molecular basis for the behavioral rescue observed (**Figure 6D**). Notably, our data show that the increased population of IMN GABA neurons persists in HOM-FlpO mice, indicating that FlpO-mediated *Scn2a* gene restoration does not reverse cellular composition (**Figure 6C**). Instead, the rescue effects may be mediated through gene expression alterations within existing cell types (**Figure 6E**). Restored expression of genes involved in axon guidance and neurodevelopment, such as *Neurod6* and *Sema3c* (70, 71), may play a critical role in re-establishing functional neural networks and ameliorating autism-related behaviors.

Compared to bulk RNA sequencing, single-cell transcriptomic approaches have provided deeper insights into the transcriptional alterations underlying ASD in mouse models carrying mutations in ASD risk genes such as *Mef2c, Myt1l,* and *Setd1a* (72–74). For instance, snRNA-seq analysis of the mPFC in mice with GABAergic neuron-specific *Mef2c* conditional heterozygosity (*Mef2c*-cHET) revealed that Pvalb-GABA neurons exhibited differential expression of genes enriched in axon development and synaptic signaling pathways (72). Similarly, large-scale snRNA-seq conducted at embryonic day 14 (E14), postnatal day 1 (P1), and day 21 (P21) in *Myt1l-*deficient mice demonstrated impaired gene programs critical for neuronal maturation and disrupted neuronal proportions in the forebrain (73). In another study, single-cell RNA-seq of adult *Setd1a* haploinsufficient (*Setd1a^+/–^*) mice revealed pronounced gene expression changes in ion transport and neurotransmission regulation within cortical neurons of the PFC. For instance, snRNA-seq analysis of the mPFC in mice with GABAergic neuron-specific *Mef2c* conditional heterozygosity (*Mef2c*-cHET) revealed that Pvalb-GABA neurons exhibited differential expression of genes enriched in axon development and synaptic signaling pathways (72). Similarly, large-scale snRNA-seq conducted at embryonic day 14 (E14), postnatal day 1 (P1), and day 21 (P21) in *Myt1l-*deficient mice demonstrated impaired gene programs critical for neuronal maturation and disrupted neuronal proportions in the forebrain (73). In another study, single-cell RNA-seq of adult *Setd1a* haploinsufficient (*Setd1a^+/–^*) mice revealed pronounced gene expression changes in ion transport and neurotransmission regulation within cortical neurons of the PFC (74). Furthermore, an integrative single-cell transcriptomic analysis of multiple ASD risk gene mutant models identified convergent gene expression changes, including alterations in the solute carrier family of transporters across diverse cell types (75). Consistent with this, our ClueGO analysis also revealed enrichment of amino acid transport ontologies. Together, these findings highlight neuronal subtype-specific differences across diverse cell types (75). Consistent with this, our ClueGO analysis also revealed enrichment of amino acid transport ontologies. Together, these findings highlight neuronal subtype-specific vulnerabilities and gene regulatory disruptions induced by ASD risk genes, suggesting their critical roles in fundamental transport processes in the brain.

We observed that the IMN GABA neurons persist as a distinct population in the mPFC of *Scn2a*-deficient mice, suggesting that these cells may represent a terminal fate rather than a transient stage destined for further differentiation. While IMN GABA neurons are considered immature due to their transcriptional features, recent comprehensive transcriptomic analyses have shown that they migrate from the subventricular zone to the olfactory bulb and may serve granule cells and periglomerular cells rather than progressing into other GABAergic subtypes (49, 76, 77). Our findings raise the possibility that *Scn2a* deficiency may interfere with this migratory or maturation process, leading to the retention of IMN GABA neurons in inappropriate brain regions. Further longitudinal studies across developmental time points, as well as functional validation, are necessary to determine whether *Scn2a* loss results in disrupted migration or delayed maturation. Future directions also include applying earlier FlpO interventions to evaluate whether altered cellular populations can be restored. Additionally, FlpO-rescued genes identified in this study, such as *Htr7*, *Grin3a*, and *Kcnip4* (neurotransmitter receptor-related), and *Neurod6*, *Sema3c*, and *Sema6a* (neuron development-related), can be further validated *in vivo* to investigate their roles in the rescue mechanism. These investigations are necessary to uncover the molecular players involved in *Scn2a* restoration-induced functional rescue.

Together, our snRNA-seq results revealed a cellular proportion change caused by *Scn2a* deficiency. We demonstrated that IMN GABA neurons in the *Scn2a*-deficient mPFC had distinct transcriptional regulation in GABAergic neurons. Differentially expressed genes in both GABAergic neurons and glutamatergic neurons highlighted ontologies related to ion channel-related and synapse organization genes. *Scn2a* gene restoration impacted cellular interactions across cell types and altered gene expression in both GABAergic neurons and glutamatergic neurons. Our results provide a transcriptomic framework that can be used to investigate the loss-of-function effects of the *Scn2a* gene at the molecular level across different cell types to inform disease mechanisms for intervention.

## Author’s contribution

Y.-E.Y., P.M. and Z.T. implemented the codes, analyzed the data, and drove the study design. Z.Z. wrote and revised the manuscript. Y.-E.Y., J.Z., X.C., M.R., M.E., B.D., and M.H. performed wet lab experiments. H.K., L.C.D., B.J., H.G., N.L., Y.L., J.K., and P.B. supervised data analyses. Y.Y. reviewed and revised the manuscript. All authors reviewed the manuscript.

## Acknowledgements

Research reported in this study was supported by National Institute of Neurological Disorders and Stroke (NINDS) of National Institute of Health under award numbers R01NS117585 and R01NS123154. This work was also supported by Hodgkin-Huxley Research Grant from FamilieSCN2A Foundation. Data analysis was supported by the Simon Comprehensive Cancer Center (Grant P30CA082709), Purdue University Institute for Cancer Research (Grant P30CA023168) and the Walther Cancer Foundation.

## Supplementary Figure Legends

**Supplementary figure 1.**
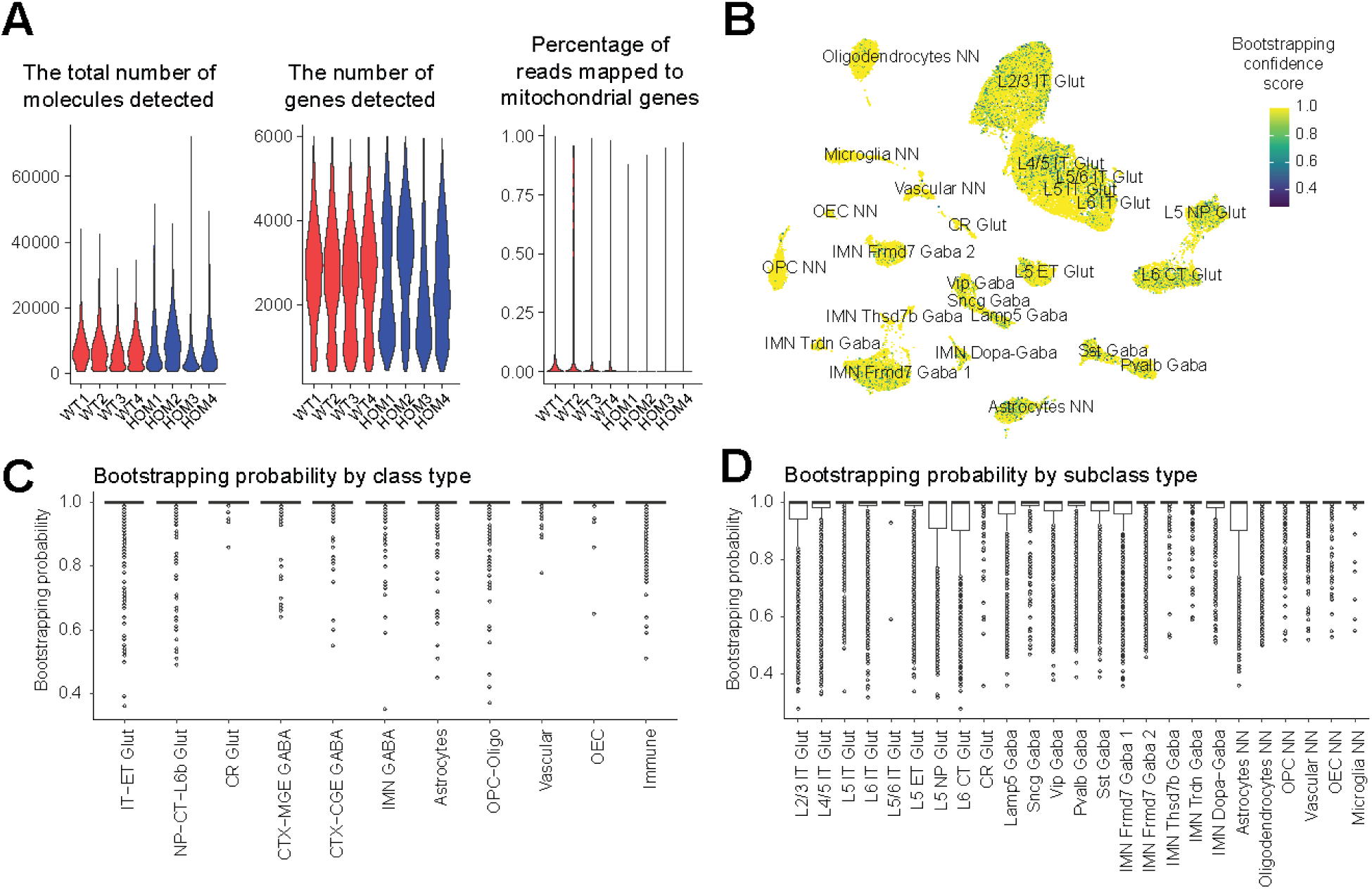
Quality control metrics of single-nucleus RNA-sequencing dataset and MapMyCells bootstrapping confidence scores for cell annotations (A) Violin plots showing the distribution of quality control metrics for each sample, including the total number of molecules, the number of detected genes, and the percentage of reads mapped to mitochondrial genes. (B) UMAP showing cell distribution represented by bootstrapping confidence score across subclass types. Note that the scores close to 1 indicate robust cell annotation. (C) Bootstrapping probability per cell across each class demonstrated that most cells showed high probability with 1. (D) Bootstrapping probability per cell across each subclass demonstrated that most cells showed high probability with 1.

**Supplementary Figure 2.**
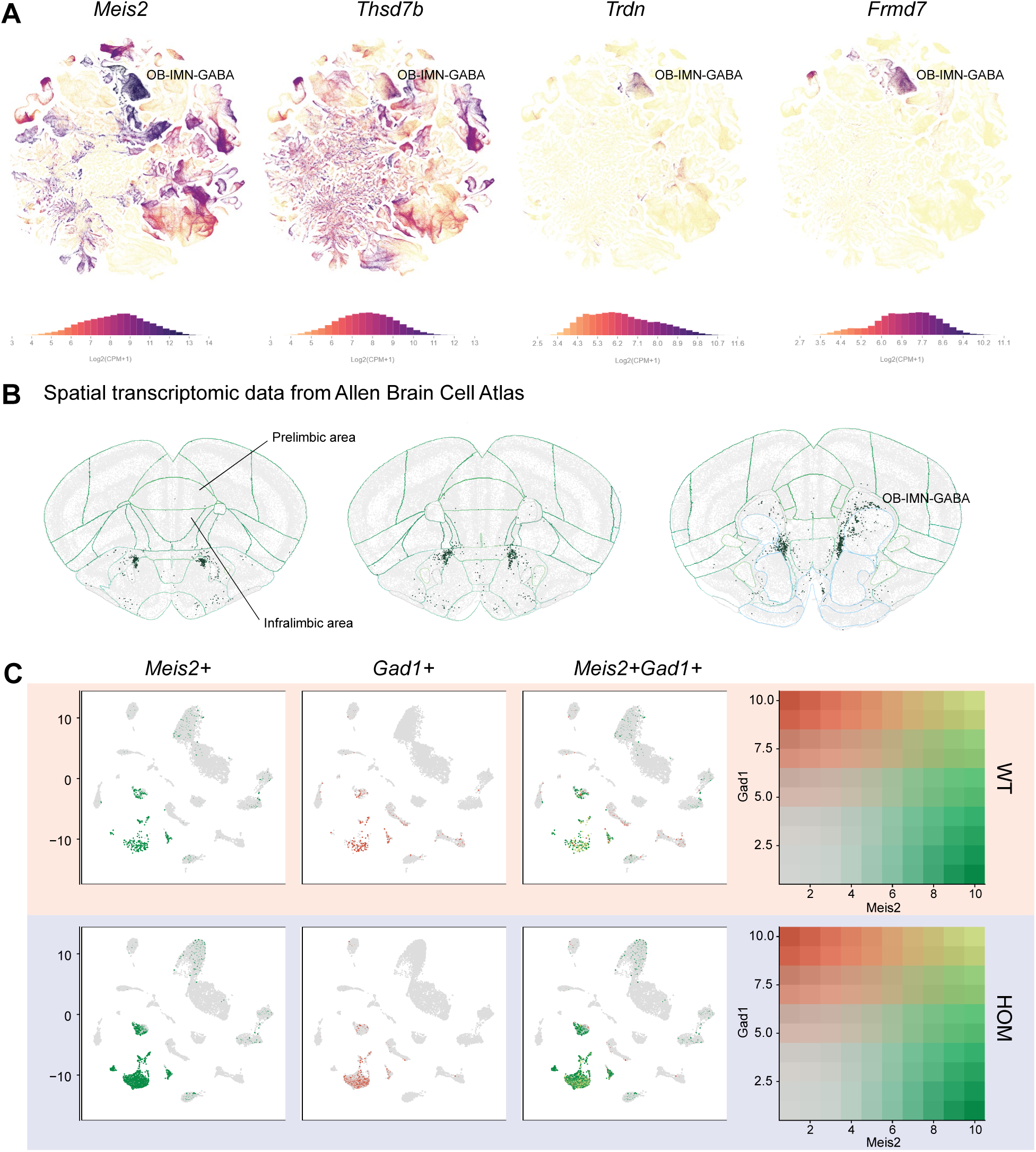
Canonical OB-IMN-GABA marker genes are enriched in IMN GABA neurons of the mPFC. (A) Expression of canonical marker genes for IMN GABA subtype visualized in the UMAP embedding of the whole brain single-cell RNA-seq dataset from the Allen Brain Cell Atlas. *Meis2* is a shared marker gene for the OB-IMN-GABA class and five IMN GABA subclasses. Subtype-specific marker genes include *Thsd7b* for IMN Thsd7b Gaba, *Trdn* gene for IMN Trdn Gaba, and *Frmd7* gene for IMN Frmd7 Gaba1 and IMN Frmd7 Gaba2. Each plot is colored by log-normalized gene expression. CPM, counts per million (B) Spatial distribution of OB-IMN-GABA cells across brain regions, based on spatial transcriptomic data from Allen Brain Cell Atlas (C) Feature plots demonstrating the expression of *Meis2* (Green) and *Gad1* (Red) gene in WT and HOM. *Meis2* expression is localized predominantly to IMN GABA neurons and overlaps with Gad1, a GABAergic neuron marker, appearing as yellow in the merged plots. The heatmap (right) display the co-expression of Meis2 (x-axis; green) and Gad1 (y-axis; red), with color intensity representing normalized expression levels.

**Supplementary figure 3.**
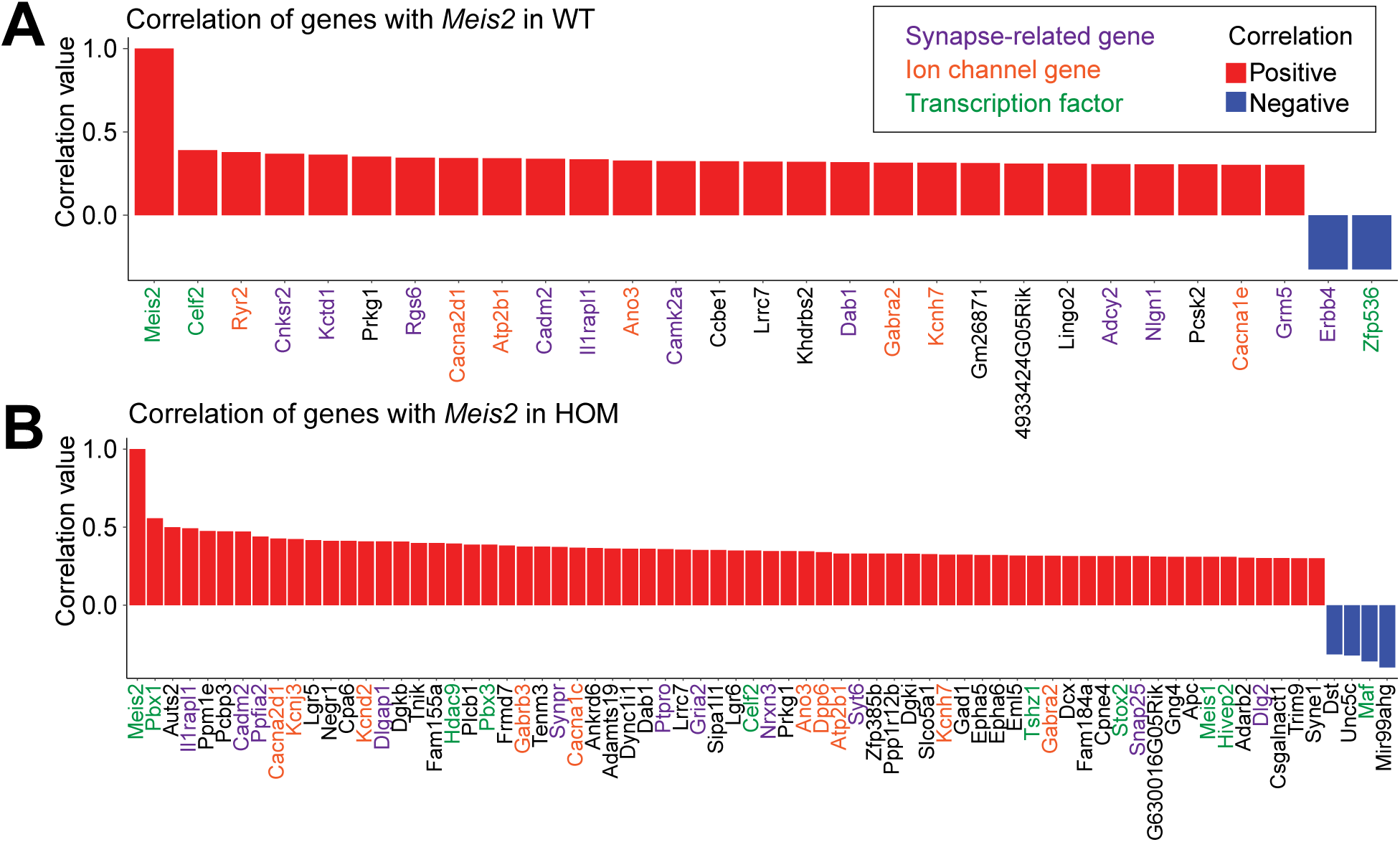
Altered *Meis2*-associated transcriptional networks in *Scn2a*- deficient mice. (A-B) Spearman correlation analysis of genes co-expressed with *Meis2* in WT (A) and HOM (B) mice. Bars represent genes with an absolute correlation value > 0.3. Red bars indicate positive correlation with *Meis2*; blue bars indicate negative correlation with *Meis2*. Synapse-related genes are labeled in purple, ion channel genes in orange, and transcription factors in green. Note that a greater number of *Meis2*-correlated genes were observed in HOM compared to WT.

**Supplementary Figure 4.**
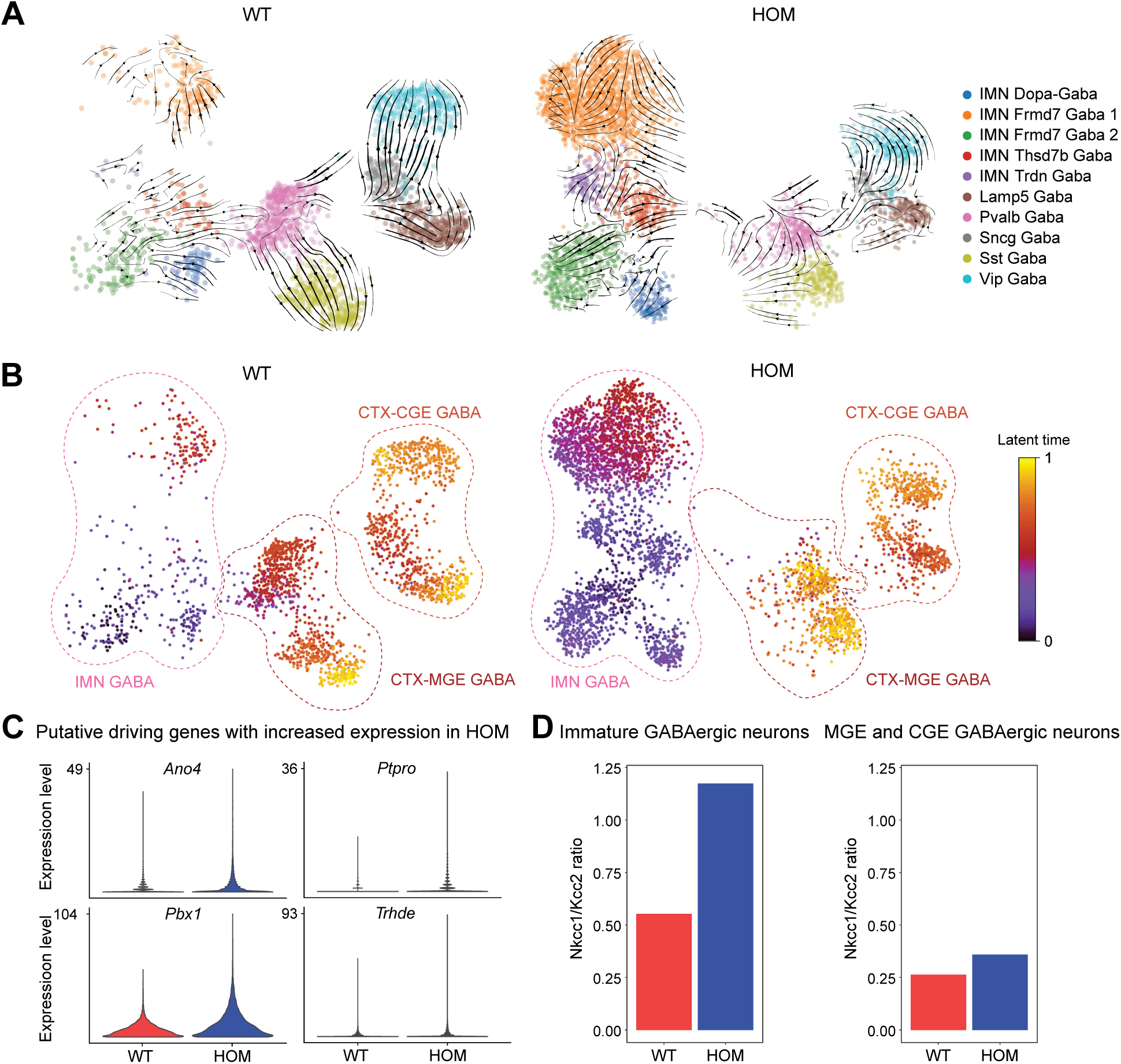
Transcriptional dynamics and cation-chloride cotransporters gene expression levels reveal immature features of IMN GABA neurons in *Scn2a*-deficient mice. (A) RNA velocity vector streams into UMAP embeddings in GABAergic neuron subtypes. Note that the IMN GABA class showed an increased population in HOM. (B) Latent time in GABAergic neurons showing transcriptomic dynamics across GABAergic cell-types in WT and HOM. (C) Violin plots showing putative driving genes contributing to the latent time in (B) with increased expression in IMN GABA neuron in HOM compared to WT. (D) Bar plot representing *Nkcc1*/*Kcc2* ratio in immature GABAergic neurons of HOM, and other GABAergic neurons (MGE and CGE GABAergic neurons).

